# A comprehensive and centralized database for exploring omics data in Autoimmune Diseases

**DOI:** 10.1101/2020.06.10.144972

**Authors:** Jordi Martorell-Marugán, Raúl López-Domínguez, Adrián García-Moreno, Daniel Toro-Domínguez, Juan Antonio Villatoro-García, Guillermo Barturen, Adoración Martín-Gómez, Kevin Troule, Gonzalo Gómez-López, Fátima Al-Shahrour, Víctor González-Rumayor, María Peña-Chilet, Joaquín Dopazo, Julio Sáez-Rodríguez, Marta E. Alarcón-Riquelme, Pedro Carmona-Sáez

## Abstract

Autoimmune diseases are heterogeneous pathologies with difficult diagnosis and few therapeutic options. In the last decade, several omics studies have provided significant insights into the molecular mechanisms of these diseases. Nevertheless, data from different cohorts and pathologies are stored independently in public repositories and a unified resource is imperative to assist researchers in this field. Here, we present ADEx (https://adex.genyo.es), a database that integrates 82 curated transcriptomics and methylation studies covering 5609 samples for some of the most common autoimmune diseases. The database provides, in an easy-to-use environment, advanced data analysis and statistical methods for exploring omics datasets, including meta-analysis, differential expression or pathway analysis.

## Background

Autoimmune diseases (ADs) are a group of complex and heterogeneous disorders characterized by immune responses to self-antigens leading to tissue damage and dysfunction in several organs. The pathogenesis of ADs is not fully understood, but both environmental and genetic factors have been linked to their development [1]. Although these disorders cause damage to different organs and their clinical outcomes vary between them, they share many risk factors and molecular mechanisms [2]. Some examples of ADs are systemic lupus erythematosus (SLE), rheumatoid arthritis (RA), Sjögren’s syndrome (SjS), systemic sclerosis (SSc), considered systemic autoimmune diseases (SADs) and type 1 diabetes (T1D), which is considered an organ-specific autoimmune disease. Most of these diseases are classified as rare given their prevalence, but altogether ADs affect up to 3 % of the population considering conservative estimates [3].

In ADs patients, the pathology is developed during several years but it is only detected when tissue damage is significant. For that reason, early diagnosis is important and complicated. Additionally, some ADs often show a non-linear outcome that alternates between active and remission stages thus making their study even more difficult. Despite huge efforts have been made to develop ADs biomarkers and therapies, these do not fit for every patient and their clinical responses differ greatly [4].

During the past decade, the use of omics technologies has provided new insights into the molecular mechanisms associated with the development of ADs, opening new scenarios for biomarkers and treatments discovery [5]. In this context, it is remarkable the characterization of the type I interferon (IFN) gene expression signature as a key factor in the pathology of some SADs, especially in SLE and SjS [6], which has improved our knowledge of the underlying molecular mechanisms and has opened new therapeutic strategies based on blocking the pathways related to this signature.

Regardless of the large amount of omics studies describing new biomarkers and therapeutic strategies in ADs [7–10], in most cases these biomarkers are not consistent across different studies or have not fully accomplished their diagnostic goals. Indeed, the widely studied IFN signature is highly variable between patients [11] and it is associated with differences in response to treatments which target it, as has been reported for example in the phase-II results of Sifalimumab clinical trial for SLE patients [12]. In addition, in most of the cases, biomarkers are defined from the analysis of a single type of omic data (commonly gene expression), but multi-omics data integration can provide a more complete understanding of molecular mechanisms and more robust and biologically relevant biomarkers.

Most of the omics datasets generated from different cohorts and studies in ADs published to date have been deposited and are available in public repositories such as Gene Expression Omnibus (GEO) [13] or ArrayExpress [14]. Although all these valuable data can be used in retrospective analyses in order to generate new knowledge and accelerate drug discovery and diagnosis, it is not easy to compare neither to integrate available data because they are generated from different platforms and/or processed with different analytic pipelines. In this context, there are great efforts from the bioinformatics community to develop standardized data analysis workflows and resources that facilitate data integration and reproducible analysis. For example, Lachmann et al. [15] have recently reprocessed a large collection of raw human and mouse RNA-Seq data from GEO and Sequence Read Archive (SRA) using a unified pipeline and they have developed the ARCHS4 as a resource to provide direct access to these data through a web-based user interface. Other singular projects such as The Cancer Genome Atlas (TCGA) [16] or the Genotype-Tissue Expression project (GTEx) [17] provide also large and homogeneously processed datasets for tumor samples and human tissues respectively. These unprecedented resources motivate the development of applications and data portals to help researchers gather information with the aim of improving diagnosis and treatment in multiple diseases, most notably in cancer research, where such information is actually being used in the clinical practice [18].

Despite such enormous potential, in the context of ADs there is a lack of a centralized and dedicated resource that facilitates the exploration, comparison and integration of available omics datasets. This is indeed an area in which this type of application would be tremendously beneficial, given that the low prevalence of each individual disease makes difficult the recruitment of large patients cohorts [4].

To bridge this gap, in this work we have compiled and curated most of the publicly available gene expression and methylation datasets for five ADs: SLE, RA, SjS, SSc and T1D. To this end, we have developed and applied homogeneous pipelines from raw data and we developed ADEx (Autoimmune Disease Explorer), a data portal where these processed data can be downloaded and exploited through multiple exploratory and statistical analyses. ADEx facilitates data integration and analysis to potentially improve diagnosis and treatment of ADs.

In order to demonstrate the potential, we queried the database to explore the expression pattern of IFN regulated genes across all autoimmune diseases. This analysis revealed that the IFN signature is consistent in SLE and SjS but it shows heterogeneity in RA samples. In a second analysis, we integrated all datasets in order to define a set of consistent biomarkers for each disease considering the expression data from multiple studies.

### Construction and content

We have prepared five different pipelines to process data for each platform (RNA-Seq, Affymetrix and Illumina gene expression microarrays, and Illumina methylation arrays 27K and 450K). All these workflows are written in R language and are publicly available in GENyO Bioinformatics Unit GitHub (https://github.com/GENyO-BioInformatics/ADEx_public). Figure 1 contains an overview of the different steps performed to prepare the data for ADEx application.

**Figure 1.**
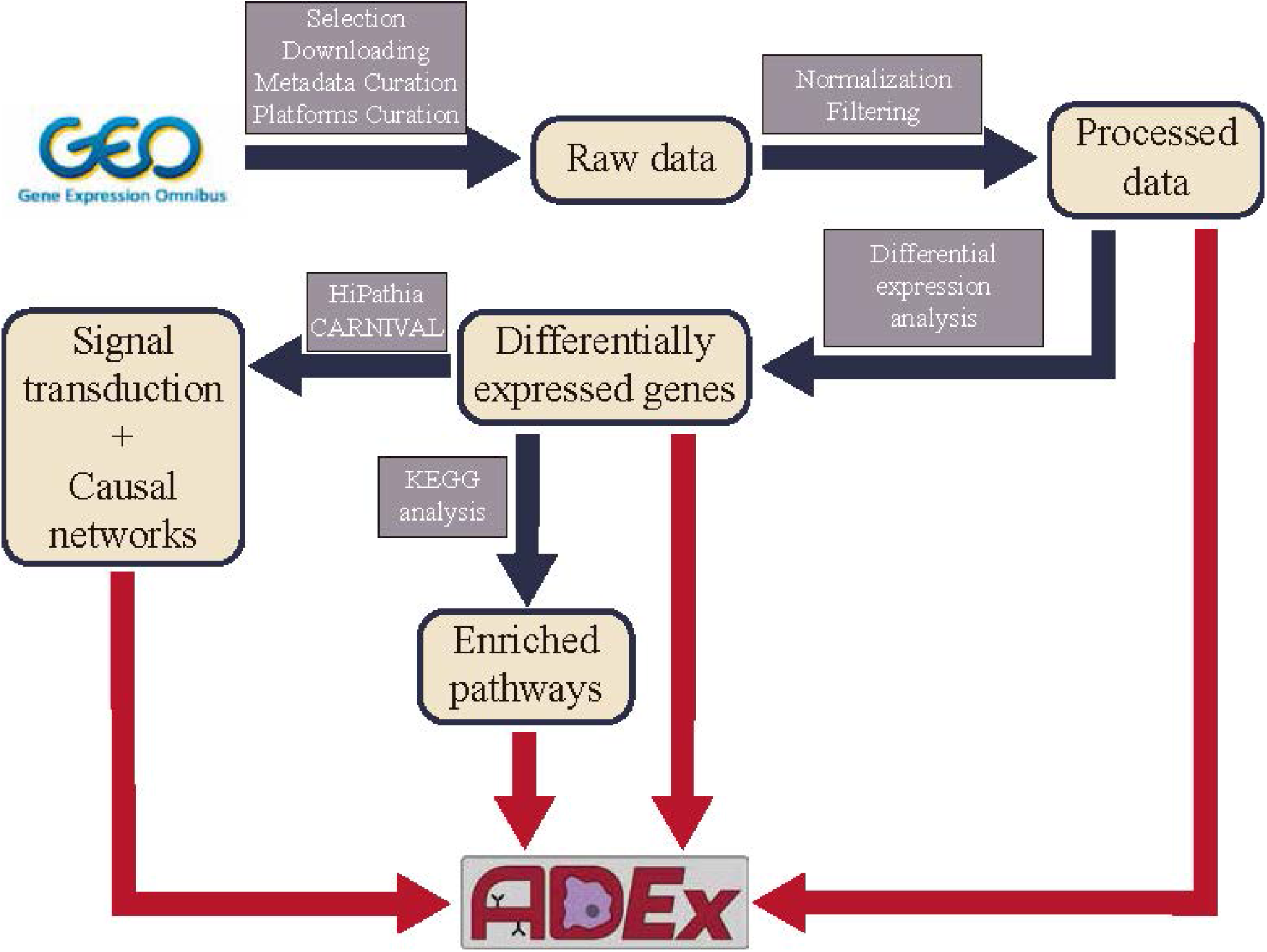
Processing pipeline for ADEx data. Black arrows indicate intermediate processing steps. Red arrows indicate the inputs to ADEx application.

### Data collection

Collection of the datasets included in ADEx was carried out by searching in the GEO web page with ADs names as key terms. We filtered the results by study type (expression profiling by array, expression profiling by high throughput sequencing and methylation profiling by array), organism (Homo sapiens) and platform manufacturer (Affymetrix or Illumina).

We downloaded the metadata for these initial datasets with GEOquery [19] R package in order to apply our inclusion criteria and exclude those studies and samples that do not meet them. We only included case-control studies from samples, which were not treated with drugs in vitro. Exclusively datasets with available raw data were considered. Studies whose controls and cases belong to different tissues were discarded. We only selected datasets with 10 samples at least. We divided the datasets containing samples from different diseases, platforms, tissues or cell types in subgroups so that these are constant and avoid possible batch effects.

82 datasets containing 5609 samples passed our filtering criteria (see Additional file 1 for complete information about all included datasets). Then we downloaded their raw data with GEOquery [19]. For expression microarrays, we downloaded CEL files and raw text files for Affymetrix and Illumina platforms respectively. For RNA-Seq, we downloaded the fastq files from the European Nucleotide Archive. For methylation microarrays, we downloaded raw methylation tables if they were available and idat files otherwise.

### Metadata curation

GEO does not require submitters to use either a fixed structure or standard vocabulary to describe the samples of an experiment. For that reason, it was necessary to manually homogenize the information provided within all the selected datasets using standardized terms. There are some methods for automatic curation of GEO metadata, but manual curation is still necessary to get high-quality metadata [20]. This metadata curation was an essential step for the following analyses and permits an easy datasets information exploration.

### Platforms curation

We have used a total of 12 different gene expression platforms from microarray and RNA-Seq technologies. Microarray platforms quantify expression levels in probes. In order to match probe identifiers to gene names, platforms annotation files are available from GEO. However, we found that some of these annotation files match probes to inappropriate gene names. On the one hand, some platforms save gene names with errors due to the conversion of gene names such as MARCH1 or SEPT1 into dates, a common error that has been reported previously [21]. In these cases, we fixed manually these genes in the annotation files. On the other hand, some platforms use obsolete or different aliases to refer to the same genes. We used human genes’ information from NCBI repository in order to match aliases with actual official gene symbols and substituted them in the platform annotations.

### Data processing

Raw data from Illumina expression microarrays were loaded by reading the plain text files. In order to remove background noise, we kept only the probes that had a Detection P-value lower than 0.05 in 10 % of the samples. Then we performed a background correction and quantile normalization [22] using neqc function from limma package [23].

CEL files from Affymetrix expression microarrays platforms were loaded to R environment with affy package [24]. To filter low intensity probes, we removed all probes with an intensity lower than 100 in at least 10 % of the samples. Normalization was carried out computing Robust Multichip Average (RMA) normalization [25] with affy package [24].

For RNA-Seq datasets, fastq files were aligned to human transcriptome reference hg38 using STAR 2.4 [26] and raw counts were obtained with RSEM v1.2.31[27] with default parameters. Raw counts were filtered using NOISeq R package [28], removing those features that have an average expression per condition lower than 0,5 counts per million (CPM) and a coefficient of variation (CV) higher than 100 in all conditions. Counts normalization was carried out with TMM method [29].

We translated microarrays probes identifiers to gene symbols using our curated annotation tables. For those genes targeted by two or more microarray probes, we calculated the median expression values of all their targeting probes. For RNA-Seq, we translated ENSEMBL identifiers to gene symbols using biomaRt package [30, 31].

Methylation raw data are available in GEO as idat or text files depending on the dataset. Idat files were read with minfi package [32], while text files were read in the R environment. In both cases, poorly performing probes with a detection P-value above 0.05 in more than 10 % of samples were removed. Probes adjacent to SNPs, located in sexual chromosomes or reported to be cross-reactive [33] were also removed. We normalized the methylation signals using quantile normalization with lumi package [34]. Finally, for datasets generated with 450k platform, we applied BMIQ normalization [35] using wateRmelon package [36] in order to correct for the two types of probes contained in this platform.

### Differential expression analysis

We performed a differential expression analysis in all datasets independently towards the identification of differential patterns among disease samples and healthy controls. These analyses were performed in different ways depending on the source of data. Gene expression profiles from microarray platforms were carried out by the standard pipeline of limma package [23]. We used lmFit function to fit a linear model to the gene expression values followed by the execution of a t-test by the empirical Bayes method for differential activity (eBayes function). On the other hand, gene expression profiles from RNA-Seq platforms were analyzed by the standard pipeline of DESeq2 package [37]. In both cases, differential expression analysis provided P-values, adjusted P-values by False Discovery Rate (FDR) and log2 Fold-Change (FC).

### Pathway analysis

Pathway enrichment analysis was precomputed for each expression dataset using differential expression analysis results. We considered DEGs those genes with a FDR lower than 0.05 and we performed hypergeometric tests to check if each pathway contains more DEGs as expected by chance. We used KEGGprofile 1.24.0 R package to perform this analysis but beforehand we manually updated its dependency, KEGG.db, the database used to perform the statistical test. The pathways were plotted using the KEGG mapper tool Search&Color Pathway, with the genes colored by their FC between case and control samples.

### Signaling network analysis

We integrated signaling network analysis applying HiPathia software [38] to gene expression data so that changes in the activity of the network from different pathways can be detected. We precomputed this analysis for each gene expression dataset. Firstly, we translated the gene expression matrix and scaled it. Then, we calculated the transduction signal and compared among conditions, cases and controls.

### Causal networks inference

We used the CARNIVAL [39] R package pipeline to analyze the causal networks architectures from gene expression data. For that aim, we followed the instructions published by their creators at https://github.com/saezlab/transcriptutorial. Briefly, differential expression analyses were performed with limma [23] and the results were used to calculate the transcription factor activities with DoRothEA [40] and the pathways activities with PROGENy [41]. These results were the input of CARNIVAL to calculate the upstream regulatory signaling pathways for each expression dataset. Finally, the results were stored in interactive html reports.

### Database architecture

Pursuing an optimal data organization and quick access to all the data in ADEx, we have enabled an internal database with PostgreSQL. We chose this technology since it is open source and it is best suited to the huge dimensionality of omics datasets.

### Webtool

ADEx user interface was designed with RStudio Shiny package. The application uses a set of external packages to perform analysis and graphics on demand. Most of the plots are generated with ggplot2 [42]. All the computations in the Meta-Analysis section are performed whenever users request them. Biomarkers analysis is performed with the Rank Products algorithm integrated in RankProd R package [43]. The tool runs on our own server with CentOS 7.0 operating system, 16 processors and 32 Gb of RAM memory.

## Utility and discussion

### Data collection and processing

ADEx contains data from 5609 samples. We have processed 82 expression and methylation datasets from case-control studies for SLE, RA, SjS, SSc and T1D diseases (see Table 1 for a summary and Additional file 1 for complete information about all included datasets). We have manually curated all metadata in order to standardize the nomenclature of phenotypes, cell types, etc. from different studies and discard samples or datasets that do not meet the selection criteria (see Construction and content section). The processed datasets are available from the Download Data section in the application.

**Table 1.** Summary of accessible studies and samples by disease and data type in ADEx.

|  | Expression | Methylation | Total |
| --- | --- | --- | --- |
| Disease | Datasets - Samples | Datasets - Samples | Datasets - Samples |
| SLE | 20 - 2053 | 13 - 628 | 33 - 2681 |
| RA | 17 - 1122 | 3 - 835 | 20 - 1957 |
| SjS | 9 - 400 | 1 - 29 | 10 - 429 |
| SSc | 5 - 229 | 1 - 37 | 6 - 266 |
| T1D | 11 - 176 | 2 - 100 | 13 - 276 |

### The ADEx application

ADEx data portal can be used to download and analyze the processed data. ADEx is freely available at https://adex.genyo.es. The tool is divided in 6 different sections arranged in different tabs (Figure 2a).

**Figure 2.**
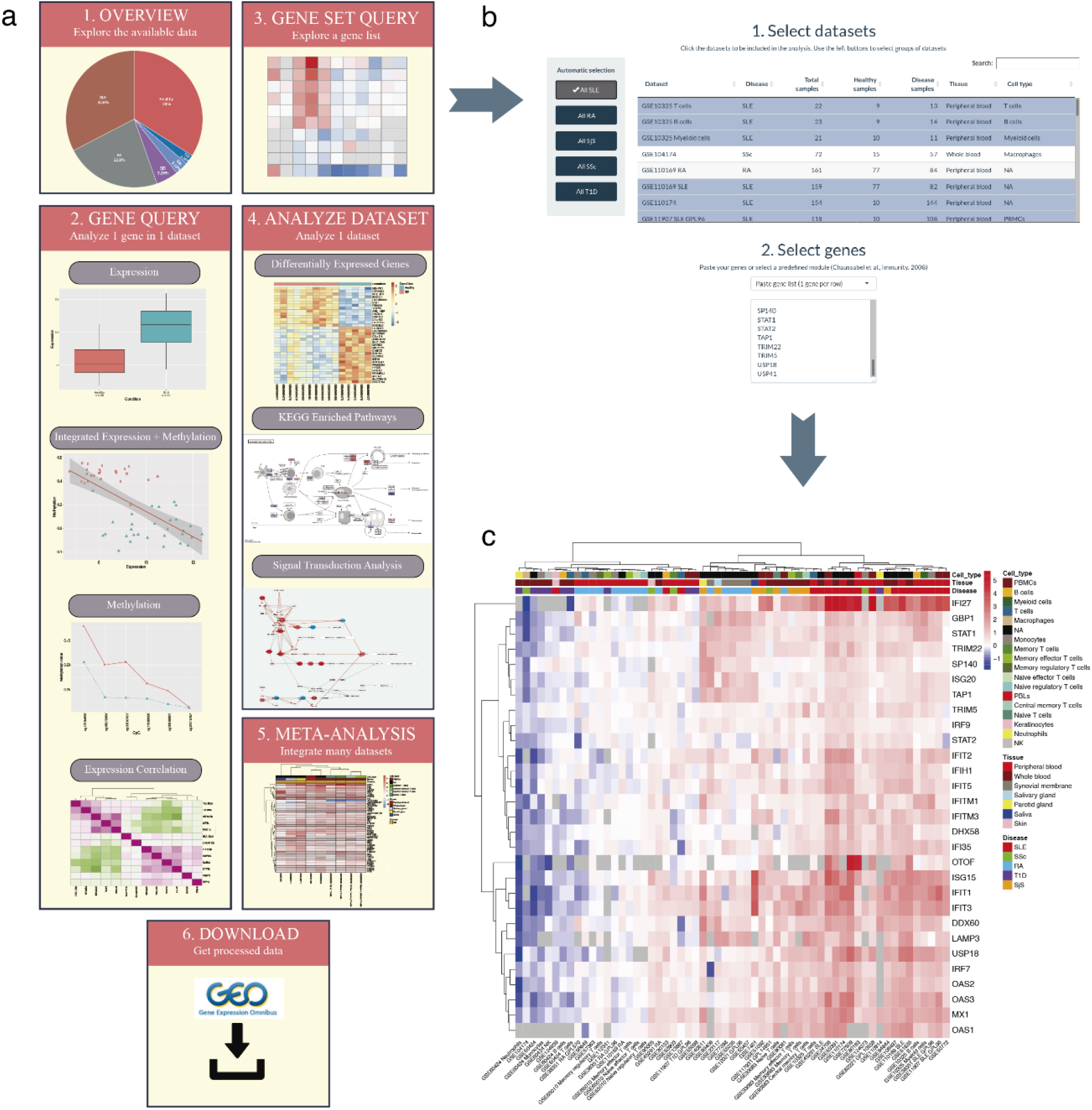
Overview of ADEx application and analysis of IFN signature across diseases. **a) ADEx has six main sections.** Section 1 provides information about available datasets. In section 2, users can explore expression and methylation for individual genes. Section 3 implements a module to explore data for a gene list, such as gene module or genes from a biological pathway, across several datasets. Section 4 allows researchers to perform analysis on individual datasets retrieving differential expression signatures and pathways and cell signaling enrichment analyses. Section 5 implements meta-analysis methods to integrate multiple datasets in order to define common biomarkers. Section 6 is for data download. **b) Gene Set Query section screenshot**. Datasets and gene set input is shown. Users select data there to plot a heatmap. **c) IFN signature expression generally separates SLE and SjS from other ADs**. Heatmap with the IFN genes generated in ADEx. Color represents the log2 FC of disease versus healthy samples (red for overexpression and blue for underexpression).

#### Section 1: Data overview

Information about the available datasets can be found in both table or pie plot formats in this section. In tables, information about the sample phenotype and their data origin is provided. In pie plots quantitative information is provided regarding the clinical and phenotype information. All this information has been extracted from GEO or from the associated published articles whenever supplied. This information can be presented individually for each dataset or grouped by disease. While a single dataset is being explored, the experiment summary is shown. Users can use this section to identify datasets of their interest to be analyzed in the following sections.

#### Section 2: Gene Query

This section was created in order to explore the expression and methylation of a specific gene, or the correlation between them, within a single dataset. Users can explore the different gene expression values for each dataset comparing case and control samples with a boxplot. Meanwhile, methylation data is presented at CpG level, so that users can select a region of the gene (e.g. promoter) and the mean methylation value for cases and controls is plotted for every CpG probe contained in the selected region.

It has been demonstrated the strong relationship of gene expression and methylation levels [44]. That is why, in this section, users can also integrate both expression and methylation values to search for direct or inverse correlations. Finally, gene expression correlation analysis can be performed in order to get insight into the relationship between different genes and to find groups of coexpressed genes.

#### Section 3: Gene Set Query

Here users can select several datasets and genes in order to explore the FC between patients and controls across studies. All datasets from a disease can be automatically selected by clicking the right buttons, or individual studies can be selected by clicking directly on the table. Users can introduce a list of genes to explore their expression, although there are several preloaded gene lists covering the coexpression modules reported by Chaussabel et al. [45]. These modules consist of sets of coexpressed genes among hundreds of samples from different diseases. Each transcriptional module is associated with different pathways and cell types, most of them related to the immune system [45]. See our use case 1 for an example of this type of analysis (Figures 2b and 2c).

#### Section 4: Analyze Dataset

In this section, we focus the analysis on whole datasets instead of individual genes. By default, a heatmap with the expression of the top 50 differentially expressed genes (DEGs) sorted by FDR is displayed. It is also possible to sort them by FC and cutoffs can be applied to both statistics. Additionally, differential expression analysis results can be downloaded as an excel table.

Furthermore, users can also study the KEGG [46] enriched pathways associated with the dataset selected. These results are precomputed using all the DEGs that have an FDR value below 0.05. A table gathers the significantly enriched KEGG pathways along with their associated hypergeometric test statistics and an interactive plot shows detailed information of the participant genes in the pathway colored according to their FC.

Beyond conventional pathway enrichment methods, we have implemented more sophisticated mechanistic models of cell signaling activity which have demonstrated to be very sensitive in deciphering disease mechanisms [38, 47] as well as the mechanisms of action of drugs [48, 49]. To offer this functionality we have applied HiPathia software [38] to gene expression data. This method estimates changes in the activity of signaling circuits defined into different pathways. With this approach, it becomes possible to study in detail the specific signaling circuits altered in ADs within the different signaling pathways. We precomputed this analysis for each dataset and the results are available as tables and interactive reports.

Finally, in this section the results of causal pathways analyses are available. We used CARNIVAL [39] software to construct the network topologies from the gene expression datasets in order to identify upstream alterations propagated through signaling networks in autoimmune diseases.

#### Section 5: Meta-Analysis

ADEx also implements meta-analysis functionalities based on gene expression data to integrate and jointly analyze different and heterogeneous datasets. We implemented a meta-analysis approach to search for biomarkers and common gene signatures across different datasets from the same or different pathologies [50] based on the FCs of each dataset and gene. Datasets have to be selected similarly to Section 3 to launch the meta-analysis. See our use case 2 for examples of this type of analysis (Figure 3).

**Figure 3.**
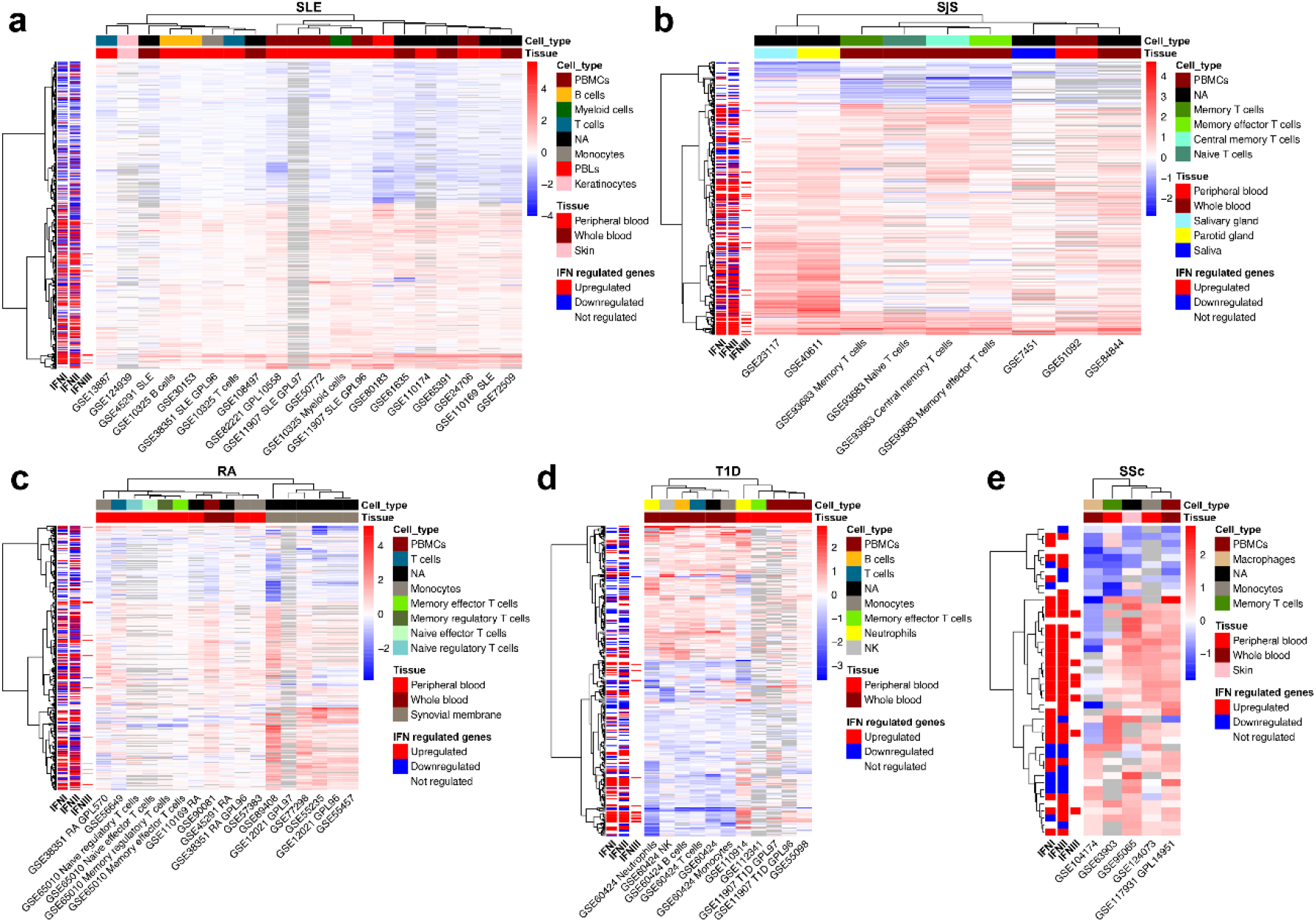
Integration of multiple datasets reveal candidate biomarkers for each disease. The observed effect of IFN I, II and III on gene expression is annotated at the left of each heatmap. Color represents the log2 FC. Heatmaps contains the significant biomarkers for **a)** SLE, **b)** SjS, **c)** RA, **d)** T1D and **e)** SSc.

#### Section 6: Download data

In this section, users can select one or several datasets and download them. Curated data is obtained with the aim of performing additional analyses externally to ADEx application.

### Use case 1: Exploring the IFN signature across diseases

Using as a query a set of genes (a gene expression signature, genes from the same pathway, etc.), it becomes straightforward to explore how the signature is expressed across different datasets or diseases. In order to demonstrate the potential of ADEx, we explored the IFN signature expression status in different diseases given its importance in the autoimmune disorders [11]. To address this goal, we evaluated the expression level across all datasets of IFN signature previously defined [51] (Figure 2b). We observed that IFN signature is strongly overexpressed in SLE and SjS patients (Figure 2c), as previously described [52, 53]. These two diseases are clearly separated from the other pathologies based on these IFN-regulated modules. RA IFN signature is highly heterogeneous, which is coherent with previous studies [54]. Interestingly, IFN modules are overexpressed in most of the RA studies that used synovial membrane tissue, while this effect is absent or very subtle in most of the RA blood studies. This is expected because the primary inflammation sites in this disease are the synovial joints [55].

### Use case 2: Biomarker discovery in ADs

To show the functionality of ADEx for biomarker discovery, we also performed a disease-centered meta-analysis with all the datasets included in the database in order to define candidate biomarkers for each disease. We removed those genes with NA values in more than 75 % of the samples and we used RankProd package [43] to calculate the Rank Product statistics and the adjusted P-value. We considered significant those genes with adjusted P-value < 0.05. Since there are datasets from different cell types, tissues, platforms and so on, our aim was to find global biomarkers independently of all those variables. We discovered 1703 consistently deregulated genes in SLE, 367 in SjS, 743 in RA, 45 in SSc and 294 in T1D (Figure 3 and Additional file 2). We used the information from Interferome database [56] to annotate each gene depending on how each type of IFN affects its expression (upregulation or downregulation). For that aim, we queried the Interferome database, searching for genes with an absolute log2 FC > 2 after IFN addition.

Given that this database contains different experimental conditions, we averaged the log2 FC and considered as genes upregulated by IFN those with an average log2 FC > 0 and as downregulated those with an average log2 FC < 0. As can be observed in Figure 3, most of SLE, SjS and RA biomarkers are expressed according to the observed IFN effect on them, supporting the major role of IFN action in these diseases. It is notable the contribution of type II IFN (IFN II) to the observed expression changes. IFN II role in ADs is frequently underestimated in favour of type I IFN (IFN I) and, in fact, IFN signature definitions commonly focus on genes regulated by IFN I [6, 10, 52]. However, it has been demonstrated that Type II IFN has a key role in ADs pathogenesis [57]. Our findings support such importance and the need to focus the attention on IFN II regulation pathways to design new therapeutic strategies.

In RA, the strongest biomarker signals come from synovial tissue studies, and these datasets are perfectly separated from the blood studies. This is coherent with the IFN signature expression results (Figure 2c).

## Conclusions

Despite that the heterogeneity of ADs is evident, there are common molecular mechanisms involved in the activation of immune responses. In this context, integrative analyses of multiple studies are crucial to discover shared and differential molecular signatures [58]. Nowadays there are many ADs datasets publicly available, but a strong computational knowledge is necessary in order to analyze them properly. With the aim of filling this gap between experimental research and computational biology, interactive easy-to-use software are valuable tools to perform exploratory and statistical analysis without strong computational expertise. This type of tool has been developed for other diseases and has helped to reuse public data and generate new knowledge and hypotheses [59–61].

A resource of this type is urged in the field of ADs to: 1) Compile available ADs’ public data in a single data portal, 2) Access to integrable data processed with uniform pipelines, and 3) Perform both individual and integrated analysis interactively. We developed ADEx database to accomplish all those objectives. Then, we used ADEx data and functions to illustrate our tool potential exploring the IFN signature in different diseases and revealing genes consistently over-and underexpressed which could be good biomarkers for these diseases.

As far as we know, ADEx is the first ADs omics database and we expect it to be a reference in this area. During the coming years, ADEx will be expanded including data from more ADs and other omics.

## Supporting information

Additioinal file 2

## Competing interests

The authors declare that they have no competing interests.

## Funding

This work is partially funded by FEDER/Junta de Andalucía-Consejería de Economía y Conocimiento (Grant CV20-36723), Consejería de Salud (Grant PI-0173-2017) and Innovative Medicines Initiative (grant GA-115565).

## Authors’ contributions

PCS conceived and directed the project. JMM designed the web functionality and interface and prepared the processing pipelines. RLD processed the data. AGM designed and implemented the SQL database and its communication with the website. DTD, KT, GGL and FA contributed to the use cases. JD, AMG and MPC implemented HiPathia analysis. JSR implemented CARNIVAL analysis. VGR, MAR, JAVG and GB tested the software and provided improvements. PCS, JMM, RLD and AGM wrote the manuscript. All authors reviewed and approved the final manuscript.

## Acknowledgements

We would like to thank all the authors of the datasets included in ADEx. We also would like to thank Alberto Ramírez for his technical support during the implementation of ADEx in our server. This work is part of the JMM’s PhD thesis. JMM is enrolled in the PhD program in Biomedicine at the University of Granada, Spain.

## Notes

https://adex.genyo.es

### Competing Interest Statement

The authors have declared no competing interest.

