## Additional file 1 for "A comprehensive and centralized database for exploring omics data in Autoimmune Diseases"

| Dataset | Studied disease | Experimental strategy | Platform | Sample size | Reference |
|---|---|---|---|---|---|
| GSE10325 | SLE | Expression profiling by array | [HG-U133A] Affymetrix Human Genome U133A Array | 67 | (Hutcheson et al., 2008) |
| GSE104174 | SSc | Expression profiling by high throughput sequencing | Illumina HiSeq 2500 (Homo sapiens) | 72 | (Moreno-Moral et al., 2018) |
| GSE108497 | SLE | Expression profiling by array | Illumina HumanHT-12 V4.0 expression beadchip | 512 | NA |
| GSE110007 | SjS | Methylation profiling by array | Illumina HumanMethylation450 BeadChip (HumanMethylation450_15017482) | 31 | (Cole et al., 2016) |
| GSE110169 | SLE, RA | Expression profiling by array | [HG-U219] Affymetrix Human Genome U219 Array | 234 | NA |
| GSE110174 | SLE | Expression profiling by array | [HT_HG-U133_Plus_PM] Affymetrix HT HG-U133+ PM Array Plate | 154 | NA |
| GSE110607 | SLE | Methylation profiling by genome tiling array | Illumina HumanMethylation450 BeadChip (HumanMethylation450_15017482) | 104 | (Ulff-Møller et al., 2018) |
| GSE110914 | T1D | Expression profiling by high throughput sequencing | Illumina HiSeq 2500 (Homo sapiens) | 42 | (Vecchio et al., 2018) |
| GSE112341 | T1D | Expression profiling by high throughput sequencing | Illumina HiSeq 2500 (Homo sapiens) | 22 | (Gao et al., 2019) |
| GSE117931 | SSc | Expression profiling by array, Methylation profiling by genome tiling array | llumina HumanHT-12 WG-DASL V4.0 R2 expression beadchip, Illumina HumanMethylation450 BeadChip (HumanMethylation450_15017482) | 74 | NA |
| GSE11907 | SLE | Expression profiling by array | [HG-U133A] Affymetrix Human Genome U133A Array [HG-U133B] Affymetrix Human Genome U133B Array | 546 | (Chaussabel et al., 2008) |
| GSE12021 | RA | Expression profiling by array | [HG-U133A] Affymetrix Human Genome U133A Array [HG-U133B] Affymetrix Human Genome U133B Array | 57 | (Huber et al., 2008) |
| GSE124073 | SSc | Expression profiling by high throughput sequencing | Illumina HiSeq 2000 (Homo sapiens) | 73 | (Mariotti et al., 2019) |
| GSE124939 | SLE | Expression profiling by high throughput sequencing | Illumina HiSeq 4000 (Homo sapiens) | 72 | (Tsoi et al., 2019) |
| GSE13887 | SLE | Expression profiling by array | [HG-U133_Plus_2] Affymetrix Human Genome U133 Plus 2.0 Array | 27 | (Fernandez et al., 2009) |
| GSE23117 | SjS | Expression profiling by array | [HG-U133_Plus_2] Affymetrix Human Genome U133 Plus 2.0 Array | 15 | (Greenwell-Wild et al., 2011) |
| GSE24706 | SLE | Expression profiling by array | Illumina HumanWG-6 v3.0 expression beadchip | 48 | (Li et al., 2011) |
| GSE27895 | SLE | Methylation profiling by array | Illumina HumanMethylation27 BeadChip (HumanMethylation27_270596_v.1.2) | 23 | (Jeffries et al., 2011) |
| GSE30153 | SLE | Expression profiling by array | [HG-U133_Plus_2] Affymetrix Human Genome U133 Plus 2.0 Array | 26 | (Garaud et al., 2011) |

| Dataset | Studied disease | Experimental strategy | Platform | Sample size | Reference |
|---|---|---|---|---|---|
| GSE38351 | SLE,RA | Expression profiling by array | [HG-U133A] Affymetrix Human Genome U133A Array [HG-U133_Plus_2] Affymetrix Human Genome U133 Plus 2.0 Array | 74 | (Smiljanovic et al., 2012) |
| GSE40611 | SjS | Expression profiling by array | [HG-U133_Plus_2] Affymetrix Human Genome U133 Plus 2.0 Array | 49 | (Horvath et al., 2012) |
| GSE42861 | RA | Methylation profiling by array | Illumina HumanMethylation450 BeadChip (HumanMethylation450_15017482) | 689 | (Liu et al., 2013) |
| GSE45291 | SLE,RA | Expression profiling by array | [HT_HG-U133_Plus_PM] Affymetrix HT HG-U133+ PM Array Plate | 805 | (Bienkowska et al., 2014) |
| GSE50772 | SLE | Expression profiling by array | [HG-U133_Plus_2] Affymetrix Human Genome U133 Plus 2.0 Array | 81 | (Kennedy et al., 2015) |
| GSE51092 | SjS | Expression profiling by array | Illumina HumanWG-6 v3.0 expression beadchip | 222 | (Lessard et al., 2013) |
| GSE55098 | T1D | Expression profiling by array | [HG-U133_Plus_2] Affymetrix Human Genome U133 Plus 2.0 Array | 22 | (Yang et al., 2015) |
| GSE55235 | RA | Expression profiling by array | [HG-U133A] Affymetrix Human Genome U133A Array | 30 | (Woetzel et al., 2014) |
| GSE55457 | RA | Expression profiling by array | [HG-U133A] Affymetrix Human Genome U133A Array | 33 | (Woetzel et al., 2014) |
| GSE56606 | T1D | Methylation profiling by array | Illumina HumanMethylation27 BeadChip (HumanMethylation27_270596_v.1.2) | 100 | (Rakyan et al., 2011) |
| GSE56649 | RA | Expression profiling by array | [HG-U133_Plus_2] Affymetrix Human Genome U133 Plus 2.0 Array | 22 | (Ye et al., 2015) |
| GSE57383 | RA | Expression profiling by array | [HT_HG-U133_Plus_PM] Affymetrix HT HG-U133+ PM Array Plate | 112 | (Rosenberg et al., 2014) |
| GSE57869 | SLE | Methylation profiling by array | Illumina HumanMethylation27 BeadChip (HumanMethylation27_270596_v.1.2) | 12 | (Hong et al., 2017) |
| GSE59250 | SLE | Methylation profiling by array | Illumina HumanMethylation450 BeadChip (HumanMethylation450_15017482) | 434 | (Absher et al., 2013) |
| GSE60424 | T1D | Expression profiling by high throughput sequencing | Illumina HiScanSQ (Homo sapiens) | 134 | (Linsley et al., 2014) |
| GSE61635 | SLE | Expression profiling by array | [HG-U133_Plus_2] Affymetrix Human Genome U133 Plus 2.0 Array | 129 | NA |
| GSE63903 | SSc | Expression profiling by array | Illumina HumanHT-12 V4.0 expression beadchip | 14 | (Ayano et al., 2015) |
| GSE65010 | RA | Expression profiling by array | [HG-U133_Plus_2] Affymetrix Human Genome U133 Plus 2.0 Array | 48 | (Walter et al., 2016) |
| GSE65391 | SLE | Expression profiling by array | Illumina HumanHT-12 V4.0 expression beadchip | 996 | (Banchereau et al., 2016) |

| Dataset | Studied disease | Experimental strategy | Platform | Sample size | Reference |
|---|---|---|---|---|---|
| GSE71841 | RA | Methylation profiling by array | Illumina HumanMethylation450 BeadChip (HumanMethylation450_15017482) | 24 | NA |
| GSE72509 | SLE | Expression profiling by high throughput sequencing | Illumina HiSeq 2500 (Homo sapiens) | 117 | (Hung et al., 2015) |
| GSE7451 | SjS | Expression profiling by array | [HG-U133_Plus_2] Affymetrix Human Genome U133 Plus 2.0 Array | 20 | (Hu et al., 2007) |
| GSE77298 | RA | Expression profiling by array | [HG-U133_Plus_2] Affymetrix Human Genome U133 Plus 2.0 Array | 23 | (Broeren et al., 2016) |
| GSE80183 | SLE | Expression profiling by high throughput sequencing | Illumina HiSeq 2000 (Homo sapiens) | 16 | (Rai et al., 2016) |
| GSE82221 | SLE | Expression profiling by array, Methylation profiling by genome tiling array | Illumina HumanHT-12 V4.0 expression beadchip, Illumina HumanMethylation450 BeadChip (HumanMethylation450_15017482) | 110 | NA |
| GSE84844 | SjS | Expression profiling by array | [HG-U133_Plus_2] Affymetrix Human Genome U133 Plus 2.0 Array | 60 | (Tasaki et al., 2017) |
| GSE87095 | RA | Methylation profiling by array | Illumina HumanMethylation450 BeadChip (HumanMethylation450_15017482) | 122 | (Julià et al., 2017) |
| GSE89408 | RA | Expression profiling by high throughput sequencing | Illumina HiSeq 2000 (Homo sapiens) | 218 | (Guo et al., 2017) |
| GSE90081 | RA | Expression profiling by high throughput sequencing | Illumina HiSeq 2000 (Homo sapiens) | 24 | (Shchetynsky et al., 2017) |
| GSE93683 | SjS | Expression profiling by array | [HG-U133_Plus_2] Affymetrix Human Genome U133 Plus 2.0 Array | 48 | (Tasaki et al., 2017) |
| GSE95065 | SSc | Expression profiling by array | [HG-U133A_2] Affymetrix Human Genome U133A 2.0 Array (HGU133A2 Hs ENTREZG 19.0.0) | 33 | NA |
| GSE10325 | SLE | Expression profiling by array | [HG-U133A] Affymetrix Human Genome U133A Array | 67 | |
